# The iron-dopamine D1 coupling modulates neural signatures of working memory across adult lifespan

**DOI:** 10.1101/2023.02.09.527840

**Authors:** Jonatan Gustavsson, Jarkko Johansson, Farshad Falahati, Micael Andersson, Goran Papenberg, Bárbara Avelar-Pereira, Lars Bäckman, Grégoria Kalpouzos, Alireza Salami

## Abstract

Brain iron overload and decreased integrity of the dopaminergic system have been independently reported as brain substrates of cognitive decline in aging. Dopamine (DA), and iron are co-localized in high concentrations in the striatum and prefrontal cortex (PFC), but follow opposing age-related trajectories across the lifespan. DA contributes to cellular iron homeostasis and the activation of D1-like DA receptors (D1DR) alleviates oxidative stress-induced inflammatory responses, suggesting a mutual interaction between these two fundamental components. Still, a direct in-vivo study testing the iron-D1DR relationship and their interactions on brain function and cognition across the lifespan is rare. Using PET and MRI data from the DyNAMiC study (n=180, age=20-79, %50 female), we showed that elevated iron content was related to lower D1DRs in DLPFC, but not in striatum, suggesting that dopamine-rich regions are less susceptible to elevated iron. Critically, older individuals with elevated iron and lower D1DR exhibited less frontoparietal activations during the most demanding task, which in turn was related to poorer working-memory performance. Together, our findings suggest that the combination of elevated iron load and reduced D1DR contribute to disturbed PFC-related circuits in older age, and thus may be targeted as two modifiable factors for future intervention.

**Highlights:**

- First study demonstrating the association between regional iron and dopamine D1DR in adult humans.
- The interplay between age-related elevated iron and diminished D1DR explained lower task-related brain activity, which in turn was related to poorer task performance.
- Our findings iron-DA coupling can help progress the understanding of the mechanisms behind DA-related neurodegeneration.

## 1 Introduction

Aging is associated with cognitive decline (Gorbach et al., 2017; Salthouse, 2012) and concomitant alterations in structural, functional, and molecular properties of the brain (Andrews-Hanna et al., 2007; Bäckman et al., 2000; Gorbach et al., 2017; Zimmerman et al., 2006). Elevated iron content and decreased integrity of the dopaminergic system are typically reported as independent brain substrates of age-related cognitive decline (Bäckman et al., 2006, 2011; Cools & D’Esposito, 2011; Daugherty et al., 2015; Gustavsson et al., 2022; Hallgren & Sourander, 1958; Kalpouzos, 2018; Landau et al., 2009; Nyberg, Andersson, et al., 2009; Salami et al., 2019). Intracellular non-heme iron is involved in several fundamental neurobiological processes, including dopamine (DA) metabolism (Hare & Double, 2016; Mills et al., 2010). More specifically, iron is a cofactor of tyrosine hydroxylase during DA synthesis, indicating a critical role of iron in DA metabolism as well as in the development of the dopaminergic system (Erikson et al., 2001; Ortega et al., 2007; Unger et al., 2014; Zucca et al., 2017).

Iron is stored in the ferritin protein which keeps it from harming brain cells and is released upon metabolic demand. However, unbound iron accumulates with advancing aging and exerts detrimental effects on cellular integrity. Elevated iron content contributes to poorer myelin integrity (Steiger et al., 2016) and grey-matter atrophy (Daugherty & Raz, 2015, 2016), as well as altered frontostriatal activity (Kalpouzos et al., 2017; Rodrigue et al., 2020; Salami et al., 2021) along with disrupted functional connectivity in aging (Salami et al., 2018). Similar to consequences of elevated iron in aging, disruption of the DA system contributes to age-related neurocognitive impairment (Bäckman et al., 2006, 2011; Cools & D’Esposito, 2011; Landau et al., 2009; Nyberg, Dahlin, et al., 2009; Rieckmann et al., 2011; Salami et al., 2019). Given DA and iron are co-localized in high concentrations in the striatum, age-related iron dyshomeostasis combined with disturbance of the DA system may become harmful for brain integrity and cognition. Direct in-vivo studies testing the synergistic effects between the dopaminergic system and iron on brain function and cognition in aging are rare.

Past studies suggest that iron may cause neurotoxicity through DA oxidation, which may in turn contribute to loss of dopaminergic neurons (Hald & Lotharius, 2005; Hare & Double, 2016; Youdim et al., 1993; Zucca et al., 2017). Animal studies have demonstrated that iron-induced damage causing DA depletion (Poetini et al., 2018) and deterioration of dopaminergic cells (Kaur et al., 2003) could be restored after iron chelation. In contrast, a longitudinal study showed that dopaminergic cell death precedes iron elevation in parkinsonian monkey (He et al., 2003). A study with young, middle-aged, and older rhesus monkeys reported an association between iron and stimulus-evoked levels of DA (Cass et al., 2007), with older monkeys exhibiting more iron accumulation and less DA release. Although it remains unclear whether elevated iron is the cause or consequence of DA degeneration, it is relatively well acknowledged in animal studies that these two key chemical components of the brain interact with each other (Hare & Double, 2016).

As opposed to ample animal studies on iron-DA coupling, evidence for such an association across the adult lifespan in humans is scarce. A recent longitudinal study showed that developmental changes in pre-synaptic, but not post-synaptic, DA receptors were positively associated with iron accumulation through adolescence and young adulthood (Larsen et al., 2020). Still, iron and DA play ambivalent roles during early and later adulthood (Johansson et al., 2022; Kalpouzos et al., 2017; Salami et al., 2021). Recent work showed that older individuals with genetic predisposition for lower DA (by proxy of the *COMT* Val158Met genetic polymorphism) accumulated more iron in the dorsolateral prefrontal cortex (DLPFC) and striatum over 3 years (Gustavsson et al., 2022). Postsynaptic DA markers, particularly DA D1-like receptor (D1DRs) which are the most abundantly expressed receptor subtypes, are among the most age-sensitive DA systems (Karrer et al., 2019). Moreover, the activation of D1 and D2 receptors alleviates oxidative stress-induced inflammatory responses (Shao et al., 2013; Yan et al., 2015; Zhu et al., 2018). Hence, with diminished DADRs, the capacity of the protective response counteracting oxidative stress and neuroinflammation due to iron overload is lessened, and may lead to increased damage and ferroptosis (Hald & Lotharius, 2005).

Motivated by the lack of human studies, we investigated the relationship between dopaminergic receptors and iron content, and their interactions on brain function and cognition. Using the largest D1DR dataset across the world, we studied 180 healthy individuals (20-79 years old) who underwent D1DR Positron Emission Tomography (PET) assessment using [^11^C]SCH23390, iron approximation made with magnetic resonance imaging (MRI) based quantitative susceptibility mapping (QSM; (Langkammer et al., 2012)), and functional magnetic resonance imaging (fMRI) while performing a working-memory task. We first examined regional variation in the link between D1DR and iron content with the hypothesis that greater iron content is related to lower D1DR, with a possible interaction with age (C.f. Salami et al., 2021; Kalpouzos et al.,2018). We targeted DLPFC and striatum, because of abundantly expressed D1DR (Shohamy & Adcock, 2010), pronounced age-related D1DR differences (Karrer et al., 2017), and association between *COMT* polymorphism (i.e., a dopaminergic gene) and iron accumulation reported in these regions (Gustavsson et al., 2022). Given the association of iron and DA to neural circuits of working memory (c.f., Salami et al., 2021; Salami et al., 2019), we predicted an interactive effect of iron and D1DR on load-dependent BOLD modulation and working memory processing (Gustavsson et al., 2022; Spence et al., 2020). To this end, we applied multivariate partial-least squares (PLS; ((McIntosh & Lobaugh, 2004)) which enables a simultaneous analysis of iron content, D1DR, age, on BOLD associations across all task conditions in a data-driven fashion. If these variables are simultaneously related to neural circuits of WM, PLS should reveal a single network showing that older individuals with elevated iron and decreased D1DR (i.e., toxic iron-DA coupling) exhibit lower task-related BOLD-response, particularly within the frontoparietal network. However, if D1DR and iron differentially modulate BOLD response (c.f., Salami et al., 2019; Salami et al., 2021), PLS should reveal different networks.

## 2 Materials and methods

This study uses data from the DopamiNe Age Connectome Cognition (DyNAMiC) project approved by the Umeå University Regional Ethical Board. All participants signed informed consent prior to data collection (for details about DyNAMiC, see Nordin et al., 2022).

### 2.1 Participants

One-hundred and eighty adults (mean age 49.8 ± 17.4 years; range 20-79; 90 females) from Umeå, Sweden, were recruited to participate in the project and underwent the full protocol, including cognitive testing, MRI, and PET assessments. All participants were native Swedish speakers, right-handed, and had no history of neurological illnesses. Of the sample of 180 participants, 3 dropped out of PET scanning, 4 were excluded due to incidental brain abnormalities, 3 due to failed reconstruction of QSM images. Additionally, brain and behavioural data were screened for univariate and multivariate outliers. Univariate outliers were defined as values greater or less than 3.29 SD from the mean (Tabachnick & Fidell, 2013) and excluded as pairwise deletions per ROIs and modality. Multivariate outliers were identified according to Mahalanobis distance (p < 0.001; Tabachnick & Fidell, 2013). As a result, five participants were identified as univariate outliers on their iron (n=3) or D1DR (n=2) values, and three were multivariate outliers. Thus, data from 162 individuals were used in the analyses involving iron content and D1D1R. Furthermore, three outliers were identified based on their online WM performance and excluded from analyses.

### 2.2 Neuroimaging acquisition and pre-processing

Participants underwent positron emission tomography on a Discovery 690 PET/CT scanner (General Electric) and MRI on a Discovery MR750 3.0 T scanner (General Electric) equipped with a 32-channel phased-array head coil at two separate occasions at Umeå University Hospital.

#### 2.2.1 PET acquisition

PET was conducted to assess whole-brain D1DR using radioligand [^11^C]SCH23390 at rest (for details see Nordin et al., 2022). The scanning session started with a low-dose CT image, followed by an intravenous bolus injection of the radioligand. Participants were instructed to lay still and stay awake with eyes open. An individually fitted thermoplastic mask was attached to the bed surface during scanning to minimize head movement.

#### 2.2.2 Structural MR acquisition

High-resolution anatomical T1-weighted images were acquired using 3D fast-spoiled gradient echo sequence with the following parameters: 176 sagittal slices, slice thickness = 1 mm, voxel size 0.49 x 0.49 x 1 mm^3^, repetition time (TR) = 8.2 ms, echo time (TE) = 3.2 ms, flip angle = 12°, field of view (FOV) = 250 x 250 mm, no spacing.

#### 2.2.3 Iron acquisition

Images for iron estimation was obtained using a 3D multi-echo gradient-recalled echo sequence (meGRE). The parameters were as follows: 124 axial slices, voxel size 1 x 1 x 1.2 mm^3^, TR = 31 ms, flip angle = 17°, FOV = 256 x 256 mm, no spacing. The first TE was 1.78 ms followed by seven additional TEs with a 2.87 ms interval.

#### 2.2.4 Functional MR acquisition

The functional images were sampled using single-shot multiband EPI sequence with 37 axial slices, voxel size 1.95 x 1.95 x 3.9 mm^3^, 0.5 mm spacing, TR = 2.000 ms, TE = 30 ms, flip angle = 80°, and FOV = 250 × 250 mm. Ten dummy scans were collected at the start of the sequence.

#### 2.2.5 In-scanner working memory task

A numerical N-back task was administered in the scanner (Salami et al., 2018). A sequence of single numbers appeared on the screen for 1.5s, with an interstimulus interval of 0.5s. During every item presentation, subjects indicated whether the digit on the screen was the same as the one shown 1, 2, or 3 digits back by pressing one of the two adjacent buttons with the index (Yes response) or middle finger (No response). Each working-memory load condition (1-, 2-, and 3-back) was presented over nine blocks in random order (interblock interval, 22 s) with each block consisting of 10 trials. For every block, 10 trials were performed with four matches (requiring a “yes” response) and six nonmatches (requiring a “no” response). The N-back blocks were counterbalanced and trial sequence was the same for all participants. The maximum score for each condition was 81, 72, 63, respectively. Performance was calculated as an error-adjusted discrimination score by subtracting the proportion of false alarms (i.e., a wrong answer judged to be correct, or, in other terms, answering Yes, when the correct answer is No) from the proportion of correct hits (i.e., answering Yes when the correct answer is Yes).

#### 2.2.6 PET processing

PET data obtained with [^11^C]SCH23390 was processed with the following steps: To estimate receptor availability (i.e., D1DR) in targeted regions, binding potential relative to non-displaceable binding in a reference region (BP_ND_; Innis et al., 2007) was used with cerebellum as reference. The processing of the PET data included correction for head movement by using frame-to-frame image co-registration, and co-registered with T1-weighted MRI data with re-slicing to MRI voxel size using Statistical Parametric Mapping (SPM12: Wellcome Trust Centre for Neuroimaging, http://www.fil.ion.ucl.ac.uk/spm/). To model the regional time-activity course (TAC) data we used simplified reference tissue model (SRTM; Lammertsma & Hume, 1996).

#### 2.2.7 MRI processing

##### Quantitative susceptibility mapping

Processing of QSM is referred from Gustavsson et al. (2022). Approximation of iron content was inferred from susceptibility values derived from QSM images. Morphology-enabled dipole inversion (MEDI; T. Liu et al., 2011) is a method for QSM estimation that selects the solution that minimizes the discrepancy in the number of voxels belonging to edges between the susceptibility image and the magnitude image. Here, we used the recommended nonlinear variant of MEDI proposed by Liu et al. (2013). Initially, the total field map was estimated from the complex meGRE images by performing a nonlinear least square fitting on a voxel-by-voxel basis. The resulting frequency map was then spatially unwrapped using a guided region-growing unwrapping algorithm (Xu & Cumming, 1999). The background fields, the superimposed field contributions that are not caused by the sources inside the brain and mainly generated by air-tissue interference, were eliminated using a nonparametric technique based on Projection onto Dipole Fields (PDF: Liu et al., 2011). Finally, the corrected frequency map was used as input for the field-to-source inverse problem to calculate susceptibility maps. The MEDI Toolbox (http://weill.cornell.edu/mri/pages/qsm.html) was used to calculate QSM images.

Due to the singularity of dipole kernel at the centre of k-space, the generated QSM images contain relative susceptibility values, which may not necessarily be comparable across subjects. To address this issue, a region of the corticospinal tract was selected as a zero-reference region due to its resilience to age-related degeneration (de Groot et al., 2015) and stable susceptibility across adulthood (Li et al., 2014). Using white-matter areas as reference regions has previously been recommended due to their low standard deviations of susceptibility, indicating low inter-subject variation similar (Straub et al., 2017) or even lower than CSF (Deistung et al., 2013). The process of zero-referencing is described in detail by Garzón and colleagues (2017).

Automated segmentation of cortical and deep gray-matter structures was performed with the Freesurfer image analysis suite — version 6 (http://surfer.nmr.mgh.harvard.edu/) using T1-weighted images (Fischl et al., 2002; Fischl, Salat, et al., 2004; Fischl, Van Der Kouwe, et al., 2004).

Next, QSM and the segmentation results were resampled to the native structural space. Then, statistics including average and standard deviation were computed on the QSM maps. We merged segmented rostral and caudal middle frontal regions from the left and right hemispheres to form DLPFC (Fig. 1A), and the left and right caudate and putamen to form striatum (supplementary materials Fig. 1A).

**Figure 1.**
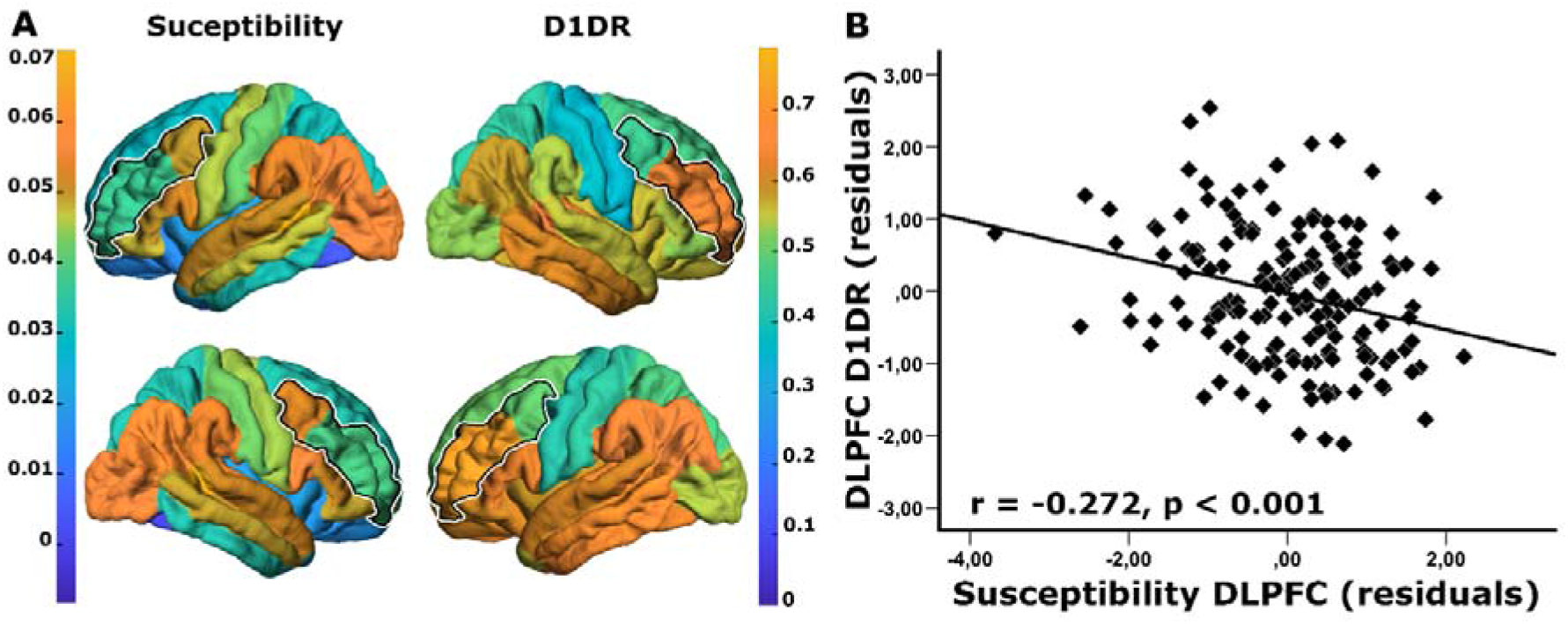
The relationship between iron content and D1DR in dorsolateral prefrontal cortex. **(A)** Surface map representing distribution of cortical iron (susceptibility in parts per million; left column) and D1DR ([^11^C]SCH23390 BP_ND_; right column) with dorsolateral prefrontal cortex (DLPFC) outlined. **(B)** Scatterplot depicting the association between greater iron content and lower D1DR in DLPFC. All values are z-transformed residuals adjusting for age.

Prior to computing statistics on the QSM maps, the boundary of segmentations was eroded by one voxel, and a fraction (15%) of the most extreme values was removed to avoid the influence of high signal from neighbouring vessels and obtain more robust estimates (Garzón et al., 2017).

#### 2.2.8 Functional MRI processing

Pre-processing of the fMRI data, performed in SPM12 software, included slice-timing correction and motion correction by unwarping and re-alignment to the first image of each volume. The fMRI volumes were then normalized to a sample-specific template generated using Diffeomorphic Anatomical Registration using Exponentiated Lie algebra (DARTEL: Ashburner, 2007), affine alignment to MNI standard space, and spatial smoothing with a 6-mm full width at half maximum (FWHM) Gaussian kernel (voxel size = 2 × 2 × 2 mm^3^).

The pre-processed fMRI data were analysed with spatiotemporal PLS (McIntosh et al., 2004; McIntosh & Lobaugh, 2004) to assess the BOLD association with iron, D1DR, and age across the three experimental WM conditions (1-, 2-, and 3-back). PLS determines time-varying distributed patterns of neural activity as a function of experimental variables (1-, 2-, and 3-back) and regional iron and D1DR. This technique allows for the identification of patterns/networks, which reflect association changes across all regions of the brain simultaneously, rather than assemblies of independent regions, thus ruling out the need for multiple-comparison correction. A detailed description of spatiotemporal PLS analysis for fMRI data has been given in previous reports (Garrett et al., 2010; Grady & Garrett, 2014; Salami et al., 2010, 2012, 2014).

The onset of each stimulus within each block of images (1-, 2-, and 3-back) was averaged across blocks for each condition. A cross-block correlation matrix was computed as the correlation between neural activity across experimental conditions (1-, 2-, and 3-back) and variables of interest (age, D1DR, iron) across different regions. Then, the correlation matrix was decomposed using singular value decomposition (SVD), to identify a set of orthogonal latent variables (LVs) representing linear combinations of the original variables:

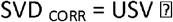

This decomposition produces a left singular vector of regional susceptibility weights (U), a right singular vector of BOLD weights (V), and a diagonal matrix of singular values (S). This analysis produces orthogonal LVs that optimally represent relations between the variables of interest (age, D1DR, iron) and BOLD. Note that PLS is not mathematically susceptible to collinearity similar to the multiple regression approach (Leibovitch et al., 1999). Each LV contains a spatial pattern exhibiting the brain regions whose activity shows the strongest simultaneous relations to the input variables. To obtain a summary measure of each participant’s expression of a particular LV pattern, subject-specific “brain scores” are computed by multiplying each voxel’s (i) weight (V) from each LV (j) by the BOLD value in that voxel for person (m), and summing over all (n) brain voxels:

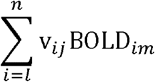

Taken together, a brain score represents the degree to which each subject contributes to the multivariate spatial pattern captured by a latent variable.

The statistical significance of each LV was assessed using permutation testing. This procedure involved reshuffling the rows of the data matrix and recalculating the LVs of the reshuffled matrix using the same SVD approach. The number of times a permuted singular value exceeds the original singular value yields the probability of significance of the original LV (McIntosh et al., 1996). In the present study, 1000 permutations were performed. In addition, the stability of voxel saliencies contributing to each LV was determined with bootstrap estimation of standard errors (SEs), using 1000 bootstrap samples (Efron & Tibshirani, 1986). The Bootstrap Ratio (BSR: the ratio between voxel saliences and estimated SEs) was computed and voxels with BSR > 3.29 (similar to a Z-score of 3.29, corresponding to p = 0.001) were considered reliable. All reliable clusters comprised contiguous voxels, with a distance between clusters of at least 10 mm. Moreover, the upper and lower percentiles of the bootstrap distribution were used to generate 95% confidence intervals (CIs) around the correlation scores to facilitate interpretation (McIntosh & Lobaugh, 2004). For instance, a significant difference between correlation scores in different conditions is indicated by non overlapping CIs. Similarly, brain or correlation scores were considered unreliable when CIs overlapped with zero.

PLS uses all conditions of an experiment and variables of interest at once, thus offering an additional dimension by simultaneously considering both similarities and differences across the experimental variables. If the variables of interest (i.e., age, iron content, D1DR) are similarly related to brain regions, PLS reveals a pattern with reliable loadings (with/without quantitative differences) for all variables. If a single variable dominates the pattern, PLS should reveal a reliable loading for that variable only. Alternatively, if different variables (e.g., D1DR and iron) differentially modulate BOLD response (e.g., Load-dependent effects of dopamine on BOLD as shown in Salami et al., (2019) versus load-independent effect of iron on BOLD shown in Salami et al., 2021), PLS may reveal two distinctive networks.

### 2.3 Additional statistical analyses

To assess age-effects on cognitive performance, a multivariate analysis of covariance (MANCOVA) was conducted with WM load-conditions as dependent variables and age-groups (younger (age 20-39) vs. middle-aged (age 40-59) vs. older (age 60-79)) as between-subjects factors. Follow-up independent t-tests were conducted to assess significant differences between age groups. To assess the relationship between iron content and D1DR, we conducted partial correlation analyses for each region of interest with iron content and D1DR as dependent variables, controlling for age. As control analyses, we performed the same analyses but included sex, education, and regional grey-matter volume as covariates.

## 3. Results

### 3.1 Demographics

Demographic information along with data on body mass index (BMI), brain volumes, D1DR, and N-back performance are presented in Table 1.

**Table 1.**
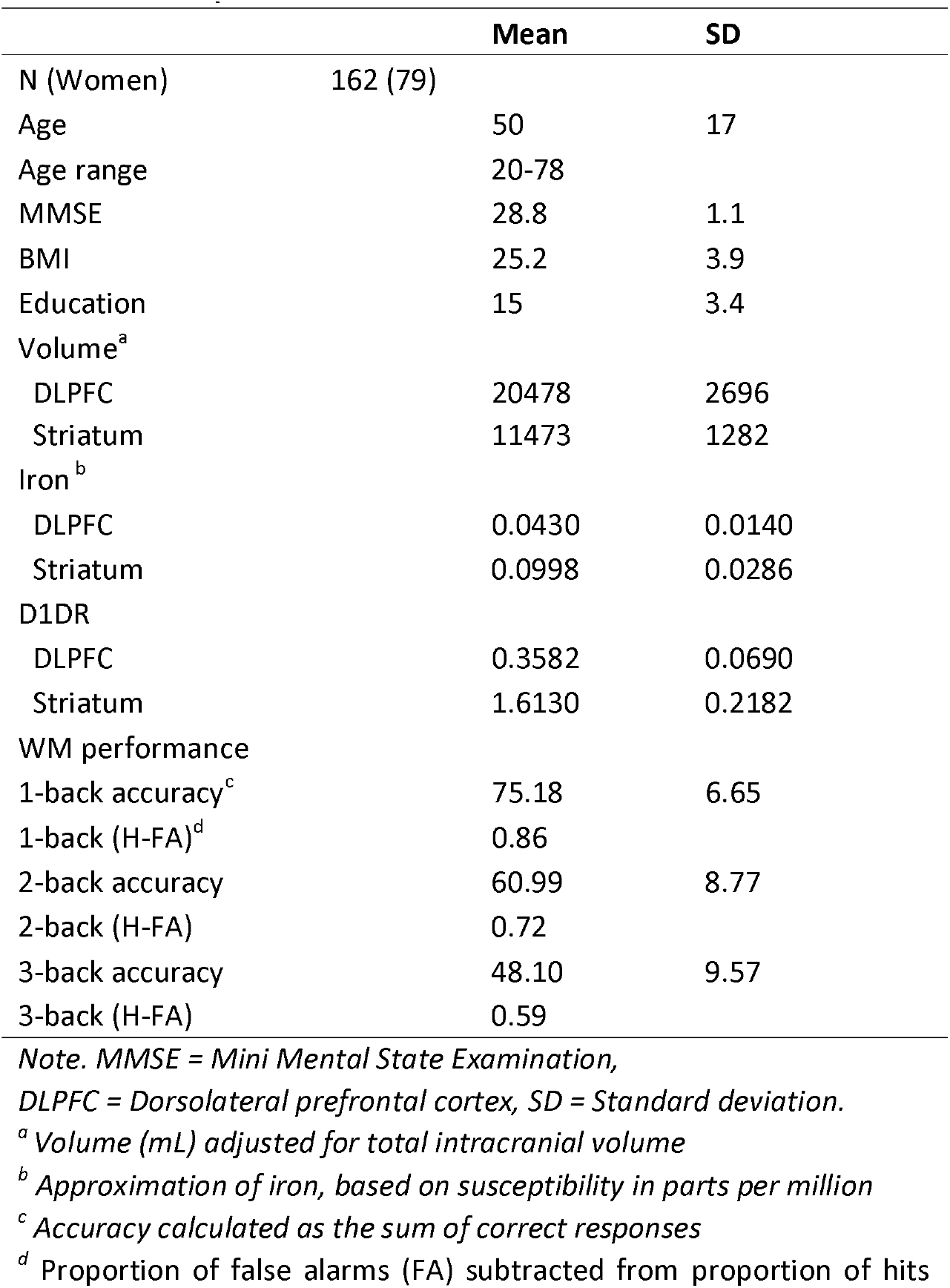
Participant characteristics.

The (H) MANCOVA conducted on N-Back performance showed a significant main effect on load conditions (F_2,155_ = 228.24, p < 0.001, Wilk’s Λ = 0.253, partial η^2^ = 0.747), and an age-group effect on load conditions (F_4,310_ = 9.43, p < 0.001, Wilk’s Λ = 0.795, partial η^2^ = 0.108). Post-hoc analysis for the lowest WM load (1-back) showed that the older participants performed less accurately compared to both middleaged (t_89_ = −3.974, p < 0.001) and younger participants (t_71_ = −5.707, p < 0.001). There was no significant difference between younger and middle-aged participants (p = 0.07). For 2-back, older participants performed significantly poorer compared to both middle-aged (t_105_ = −4.015, p < 0.001) and younger participants (t_85_ = −8.905, p < 0.001). Middle-aged participants performed significantly poorer compared to younger participants (t_85_ = −4.176, p < 0.001). Lastly, for highest WM load (3-back), older participants performed significantly poorer compared to both middle-aged (t_105_ = −5.300 p < 0.001) and younger participants (t_102_ = −7.950, p < 0.001). Middle-aged participants performed significantly poorer compared to younger participants (t_105_ = −2.921, p = 0.004).

### 3.2 Iron content in DLPFC, but not in striatum, was negatively associated with D1DR

Both striatal and DLPFC iron increased with advancing age (striatum: r = 0.551, p < 0.001; DLPFC: r = 0.244, p = 0.002). D1DRs in both regions decreased with advancing age (striatum: r = −0.62, p < 0.001; DLPFC: r = −0.565, p = <0.001).

The partial correlation analysis for iron content and D1DR in DLPFC showed a significant negative association across the whole sample (r = −0.272, p < 0.001), suggesting that greater iron content was linked to lower D1DR (Fig. 1B). Further group level analyses revealed that this correlation was similarly expressed across different age groups (Younger: r = −0.10, p = 0.48; Middle-aged: r = −0.30, p = 0.026; Older: r = −0.37, p = 0.006). However, striatal iron content was unrelated to D1DR across the whole sample (r = −0.101, p = 0.20; supplementary materials, Fig. 1B). No significant association was observed within each age groups (Ps > 0.05), except for a trend-level link in the older individuals (r = - 0.24, p = 0.08).

### 3.3 Iron-D1DR couplings in DLPFC modulates WM-related BOLD in a load-dependent fashion

Given the iron-D1DR link in DLPFC, we next investigated whether iron-D1DR coupling modulated neural correlates of PFC-related function across the adult lifespan. We used behavioural PLS to assess the presence of multivariate spatial patterns of task-related BOLD response dependent on age, iron content, and D1DR across load conditions during N-back working memory task. The analysis identified two significant latent variables (LVs). LV1 represented a brain-wide network with increased activation in older individuals in a load-independent fashion. This network was largely unrelated to D1DR or iron in DLPFC (supplementary materials 5.1.2, Table 1, Fig 2).

The second LV represented the canonical WM network (c.f. Nagel et al., 2009; Salami et al., 2018). This LV (permuted p = 0.02, 17.97% of cross-block covariance: Fig. 3A) demonstrated that older individuals with elevated iron content and lower D1DR in DLPFC exhibited lower BOLD response in the frontoparietal network during 3-back (Fig. 3B). In contrast, higher BOLD response in the frontoparietal network during 1-back was associated with older age and lower D1DR in DLPFC, but not with iron content. For 2-back, no associations were considered reliable as the CI:s overlapped with zero (Fig. 3B). As opposed to the frontoparietal WM network, the default-mode network (Fig. 3A (Blue areas); supplementary Table 2) was less deactivated during 3-back in older individuals with higher iron content and lower D1DR in DLPFC. Taken together, these results revealed that iron-D1DR coupling modulates neural correlates of the working memory in a load-dependent fashion.

**Figure 3.**
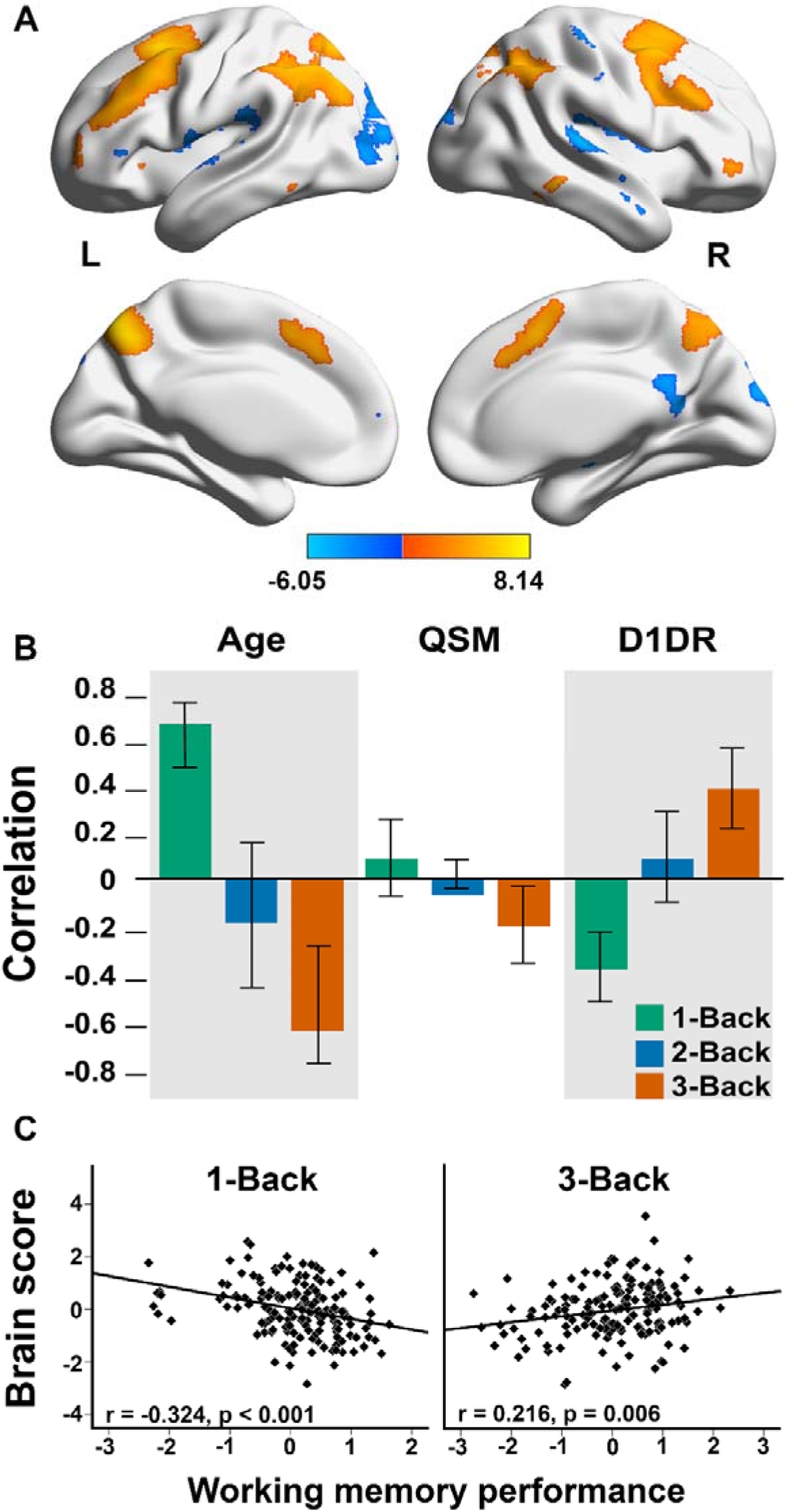
Multivariate relationship of BOLD response patterns during working-memory task identified by task PLS. A) The regions depicted in a yellow colour mainly correspond to the frontoparietal network, whereas the blue colour mainly represents the default-mode network. The frontoparietal network showed greater activation in correspondence to the behavioural measures across different task conditions, whereas the default-mode network showed greater deactivation (i.e., less activation). **B)** BOLD association across age, iron content (QSM), and D1DR. Interpretation of the figure should be as reliable positive and negative associations of activation (BOLD response) when the confidence intervals (CI:s) do not overlap with zero. As the frontoparietal network displayed increased activation, a positive association is interpreted as greater activation, whereas a negative association is interpreted as less activation. For the default-mode network the opposite applies. **C)** Greater activation was associated with poorer performance during 1-back, but greater performance during 3-back. All values are z-transformed residuals adjusting for age.

### 3.4 Load-dependent BOLD response is differentially associated with task performance

We have shown that increasing age was concomitant with increased and decreased activations within the frontoparietal network during 1-back and 3-back, respectively. We next tested the relationship of brain activation in relation to task performance. The brain score during 1-back was negatively associated with 1-back performance (r = −0.324, p < 0.001; Fig. 3C), but positively with 3- back (r = 0.216, p = 0.006; Fig. 3C). No significant association was observed during 2-back (r = 0.05, p=0.4). Taken together, our results suggest that less frontoparietal activity during 3-back as well as greater frontoparietal activity during 1-back are associated with less efficient working memory function.

### 3.5 Control analyses

To confirm that the results obtained from the partial correlation analyses were not due to confounding variables, we performed additional analyses in which we controlled for sex, education, and regional grey-matter volume. The inclusion of the additional variables did not result in a noticeable change of the outcome and all the patterns remained consistent. A full description of the results is given in the supplementary results.

## 4. Discussion

In this study, we provide first in-vivo evidence for an association between D1DR and brain iron across the adult lifespan, and how iron-D1DR synergy may contribute to disturbed brain responses during WM task performance across the lifespan. The main findings were that greater iron content was associated with lower D1DR in DLPFC, but not in striatum, across the adult lifespan. We also found that older individuals with elevated iron content and lower D1DR in DLPFC – a presumably deleterious synergy – exhibited lower frontoparietal activity along with less deactivation of the DMN during high demand condition and in turn poorer WM performance. Although the associations revealed by these data are cross-sectional, a plausible interpretation is that elevated iron and lower D1DR together form a toxic couple (c.f. Hare & Double, 2016), which would contribute to reduced network dynamics and impaired WM processing.

A key finding of the present study is the association between higher iron content and lower D1DR in DLPFC across the adult lifespan. The exact mechanism underlying the association between iron content and DA receptors is poorly understood. The dopaminergic system is important to cellular iron homeostasis, as documented in *in vitro* studies (Dichtl et al., 2018; Liu et al., 2021). Supporting this idea, we recently demonstrated that older adults with genetic predisposition for lower DA levels accumulated more iron in DLPFC and striatum over time compared to individuals with presumably higher DA levels (Gustavsson et al., 2022). Age-related differences in pre- and postsynaptic DA markers (Bäckman et al., 2006, 2010) may also contribute to iron dyshomeostasis in aging, resulting in disruption of transportation of iron via transferrin and storage in the ferritin protein (Ward et al., 2014). Disruption of iron handling may lead to increase of ferrous iron, which triggers oxidative stress, neuroinflammation, and cell loss due to ferroptosis (Daugherty et al., 2015; Mazhar et al., 2021; Salami et al., 2021). The activation of D1 and D2/3 receptors alleviates oxidative stress-induced inflammatory responses (Shao et al., 2013; Yan et al., 2015; Zhu et al., 2018). However, whether age-related loss of DA receptors leads to an iron-related increase of oxidative stress and ferroptosis, or whether an age-related increase of iron causes loss of DA receptor cells can only be determined in a longitudinal setting.

We did not observe any significant association between iron and D1DR in striatum, except for a trend-level negative association in older age. The reason for these regional variations is unclear. One possibility is that, as D1DR are more expressed in striatum compared to cortex (Shohamy & Adcock, 2010), striatal regions are less vulnerable or better at attenuating the damage from higher iron content. If too many receptors diminishes, or the threshold of the capacity for dealing with oxidative stress is exceeded, the iron-induced damage to the cells may lead to ferroptosis (Lillig et al., 2008). This concords well with Parkinson’s Disease studies reporting regional variations in synuclein pathology despite brain-wide iron accumulation (McCann et al., 2016), suggesting that additional factors may contribute to iron serving a pathological role.

The multivariate PLS analysis revealed that in older age the combination of higher iron and lower D1DR within DLPFC was related to high frontoparietal recruitment during low-demanding tasks along with weak frontoparietal upregulation at higher task demand, which in turn was associated with poorer working memory performance. These results concord with the Compensation-Related Utilization of Neural Circuits Hypothesis (CRUNCH; (Cappell et al., 2010; Mattay et al., 2006; Nyberg et al., 2014; Reuter-Lorenz & Cappell, 2008; Schneider-Garces et al., 2010)), which postulates that neural activity increases at low demanding task-levels in older adults compared to younger adults, but is reduced at more demanding levels.

The unique and shared contributions of iron content and neuroinflammation to astrocytic dysfunction in the neurovascular coupling has been proposed to account for the reduced BOLD response (Kalpouzos et al., 2017; Salami et al., 2018, 2021). Higher iron content has been related to inflammation (Haider, 2015; Salami et al., 2021), which can be detrimental to cells during prolonged periods (Hald & Lotharius, 2005; Zecca et al., 2004), thus contributing to astrocytic dysfunction. In vitro studies have demonstrated that DA receptors activation may alleviate and supress neuroinflammation (Liu et al., 2021; Shao et al., 2013; Yan et al., 2015). A more efficient dopaminergic system (e.g., manifested by greater receptor availability) protect against negative effects of iron and neuroinflammation on brain function. In support of this notion, a past study showed that age-related D1DR alteration alone may contribute to less efficient engagement of working memory circuits (Bäckman et al., 2011). Relatedly, the importance of DA for the frontoparietal BOLD response has been further demonstrated through pharmacological administration of a dopaminergic antagonist, leading to poorer working-memory performance (Fischer et al., 2010). Our results add novel contribution to the previous work, by showing that the combination of elevated iron and D1DR reduction in DLPFC in aging, possibly reflecting synergistic iron-D1DR effects, exerts a deleterious effect in neural circuits of WM. Furthermore, during the most demanding WM condition (3-back), lower activation in the frontoparietal network was related to worse performance, in concordance with previous reports (Salami et al., 2021). Longitudinal data are needed to identify the primary mechanism of disturbed working memory circuit in older age.

Our study is the first to link regional iron content to DA receptor availability, by showing that greater iron content is related to lower D1DR. Critically, an interplay between age-related elevated iron content and diminished D1 receptor availability may provide a mechanistic understanding of how iron-DA coupling may exert deleterious effects on neural function and ultimately cognition. Elevated brain iron has been implicated in several neurodegenerative disorders, including Parkinson’s disease, which is characterized by loss of dopaminergic cells (Ward et al., 2014). Thus, the observed findings have implications for better understanding the mechanisms behind DA-related neurodegeneration.

## Author credit statement

Conceptualisation: A.S. and G.K.; Software, M.A.; Methodology, J.G., G.K., G.P., A.S.; Formal Analysis J.G., G.K., A.S., J.J., F.F., G.P., M.A., and B.A.P.; Visualisation, J.G. Writing – original draft, J.G., G.K., A.S., F.F.; Writing – Review & Editing G.P., J.J., L.A., B.A.P.

## Acknowledgement

We thank the staff of the DyNAMiC project, Frida Magnusson and Emma Simonsson, staff at MRI and PET labs at Umeå University Hospital, and all our participants. We also thank Robin Pedersen for his contribution with the figures.

## Ethics approval

This study was approved by the Regional Ethical board and the local Radiation Safety Committee in Umeå, Sweden.

## Data availability statement

Data from the DyNAMiC project cannot be made publicly available due to ethical and legal restrictions. However, access to these original data may be available upon request from the corresponding author.

