## Supplemental materials for "The iron-dopamine D1 coupling modulates neural signatures of working memory across adult lifespan"

### 5. Appendix

#### 5.1 Supplementary results

##### 5.1.1 Supplementary control analyses

The MANCOVA conducted on N-Back performance showed a significant main effect on load conditions ($F_{2,153}$ = 14.556, p < 0.001, Wilk's Λ = 0.840, partial $\eta^2$ = 0.160), and an age-group effect on load conditions ($F_{4,306}$ = 8.901, p < 0.001, Wilk's Λ = 0.802, partial $\eta^2$ = 0.104). Post-hoc ANCOVA conducted on the lowest WM load (1-back) showed significant age effect on WM performance ($F_{2,159}$ = 17.07, p < 0.001, $\eta^2$ = 0.181). Older participants performed less accurately compared to both middle-aged (mean difference = -0.085, p < 0.001) and younger participants (mean difference = -0.114, p < 0.001). There was no significant difference between younger and middle-aged participants (p = 0.4). The ANCOVA for 2-back performance showed significant age effect ($F_{2,159}$ = 36.601, p < 0.001, $\eta^2$ = 0.322), as older participants performed significantly poorer compared to both middle-aged (mean difference = -0.141, p < 0.001) and younger participants (mean difference = -0.244, p < 0.001). Middle-aged participants performed significantly poorer compared to younger participants (mean difference = -0.103, p < 0.001). Lastly, the ANCOVA for the highest WM load (3-back) also showed significant age effect on performance ($F_{2,159}$ = 31.244, p < 0.001, $\eta^2$ = 0.289), older participants performed significantly poorer compared to both middle-aged (mean difference = -0.184, p < 0.001) and younger participants (mean difference = -0.284, p < 0.001). Middle-aged participants performed significantly poorer compared to younger participants (mean difference = -0.100, p = 0.017).

Iron in both striatum and DLPFC increased with age (striatum: r = 0.493, p < 0.001; DLPFC: r = 0.227, p = 0.004), whereas D1DRs decreased in both regions (striatum: r = -0.557, p < 0.001; DLPFC: r = -0.387, p < 0.001).

The associations between iron and D1DR in DLPFC and striatum remained unchanged across the whole sample and on group level (DLPFC: r = -0.290, p < 0.001; Group level: Younger: r = - 0.12, p =

0.4; Middle-aged: r = -0.32, p = 0.025; Older: r = -0.37, p = 0.007; Striatum: r = -0.091, p = 0.2; Group level: Younger: r = - 0.13, p = 0.3; Middle-aged: r = -0.09, p = 0.57; Older: r = -0.27, p = 0.055).

With regards to the relationship between WM BOLD response and WM performance, the results remained intact (1-back: r = -0.299, p < 0.001; 2-back: r = 0.089, p = 0.3; 3-back r = 0.229, p = 0.004).

#### 5.1.2 Multivariate spatial pattern results

The PLS revealed two significant LVs. LV2 exhibited a frontoparietal working memory network, and is fully described in the main text. LV1 (p < 0.001, 67.86% of cross-block covariance) identified a brain-wide network that represented regions with reduced activation (supplementary Fig. 2).

### 5.2 Supplementary figures

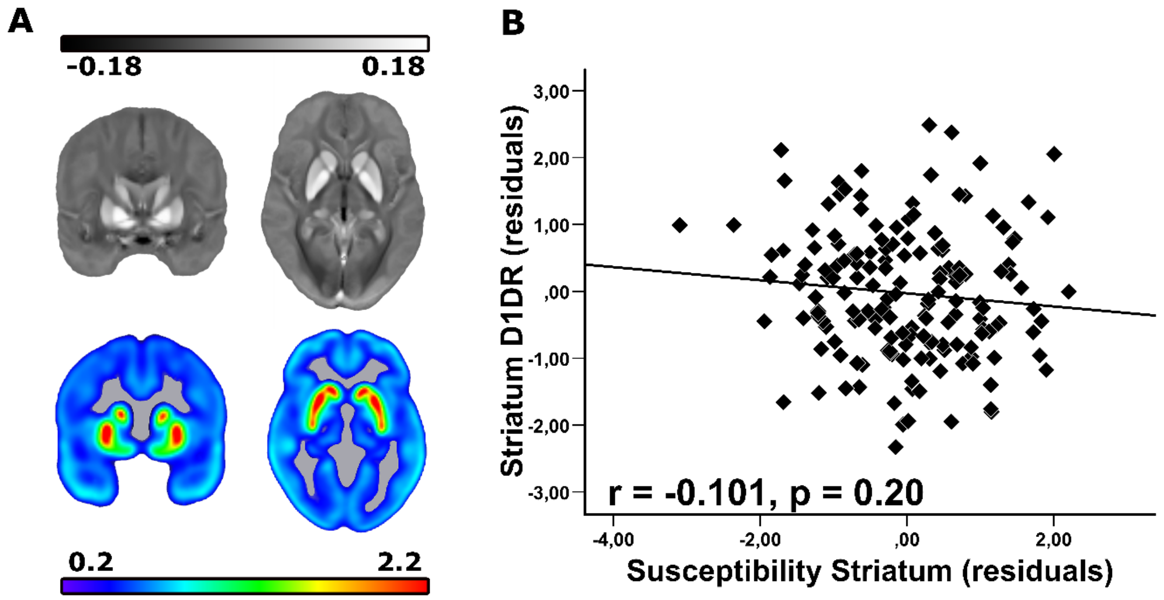


**Supplementary Figure 1** The relationship between iron content and D1DR in striatum. **(A) Top row:** QSM maps depicting average subcortical iron content (susceptibility in parts per million) with greater intensity representing higher iron content. **Bottom row:** Brain maps depicting average subcortical

D1DR availability ([$^{11}$C]SCH23390 $BP_{ND}$). **(B)** Scatterplot depicting the association between greater iron content and lower D1DR in striatum. All values are z-transformed residuals adjusting for age.

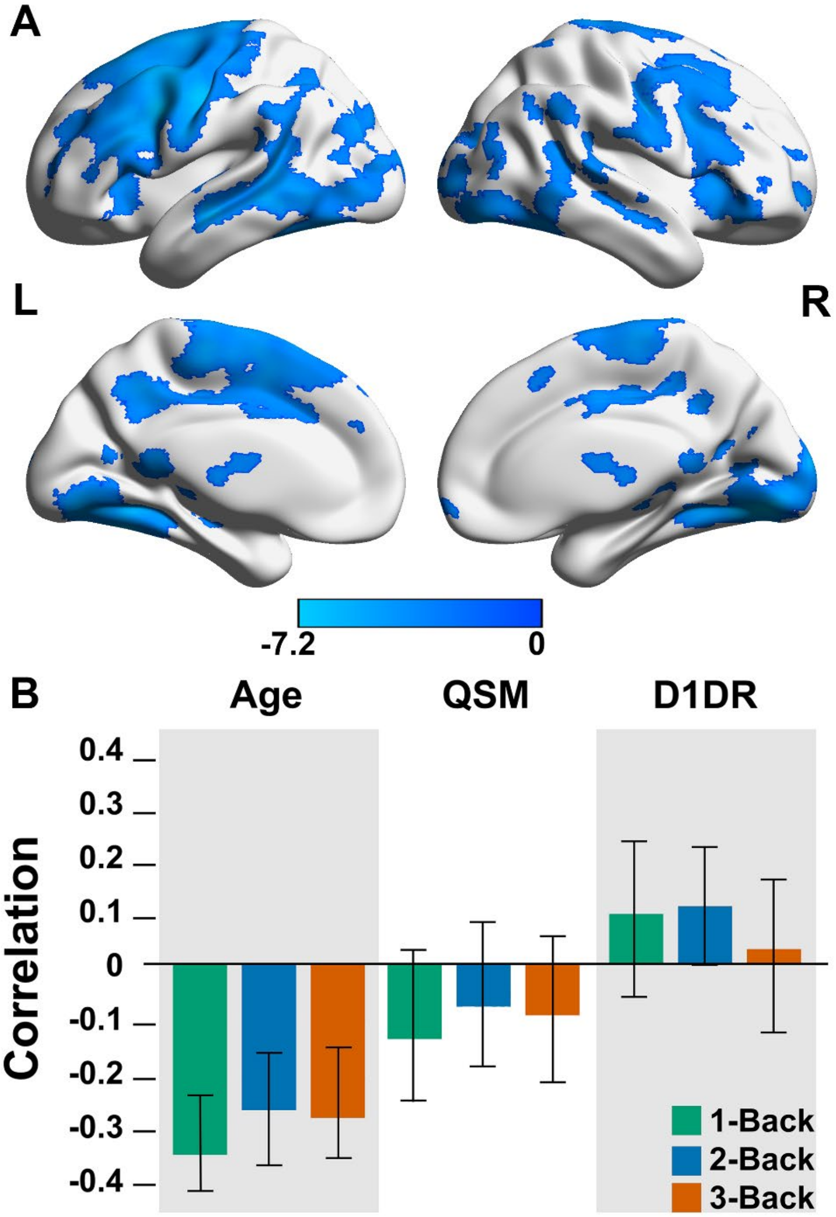


**Supplementary Figure 2.** Multivariate relationship of BOLD response patterns during working-memory task identified by task PLS. **A)** The blue coloured regions are associated with regions exhibiting lower activity during the different task conditions. **B)** Multivariate relationship of BOLD response to age, iron content (QSM), and D1DR. Interpretation of the figure should be interpreted as reliable positive and negative associations of activation (BOLD response) when the confidence intervals (CI:s) do not overlap with zero. As this network mainly showed deactivation, a positive association is interpreted as greater deactivation, whereas a negative association is interpreted as less deactivation (i.e., more activation).

#### 5.3 Supplementary Tables

**Supplementary Table 1.** Regions identified by task PLS as LV1 showing BOLD response dependent of age, iron, and D1DR. Details of the peak voxel of each cluster that resulted from thresholding by a bootstrap ratio of 3.29 (similar to a Z-score of 3.29, corresponding to p = 0.001). Only clusters that included at least 30 voxels are reported (k > 30, voxel size 2 x 2 x2 $mm^3$).

| Region | x | y | z | Bootstrapped ratio | k (cluster size) |
|---|---|---|---|---|---|
| **Decreased activity** | | | | | |
| Left lingual | 34 | 23 | 31 | -7.2113 | 15428 |
| Right precentral | 63 | 65 | 54 | -7.0954 | 15143 |
| Left precentral | 21 | 64 | 52 | -4.9573 | 2625 |
| Right caudate | 46 | 62 | 43 | -4.9304 | 1525 |
| Left postcentral | 16 | 55 | 53 | -5.5433 | 1136 |
| Left insula | 23 | 72 | 37 | -4.3837 | 946 |
| Right inferior orbitofrontal | 65 | 75 | 29 | -5.6146 | 675 |
| Right precuneus | 43 | 39 | 42 | -4.5311 | 281 |
| Left supramarginal | 11 | 37 | 54 | -4.2791 | 202 |
| Left superior orbitofrontal | 30 | 91 | 33 | -4.7799 | 201 |
| Left cerebellum | 36 | 32 | 15 | -4.1828 | 48 |
| Left hippocampus | 31 | 49 | 32 | -4.3731 | 45 |
| Left middle frontal | 26 | 90 | 44 | -3.5316 | 36 |
| Left lingual | 33 | 43 | 36 | -3.9314 | 34 |
| **Increased activity** | | | | | |
| Left cuneus | 40 | 19 | 54 | 5.9953 | 192 |

**Supplementary Table 2.** Regions identified by task PLS as LV2 showing BOLD response dependent of age, iron, and D1DR. Details of the peak voxel of each cluster that resulted from thresholding by a bootstrap ratio of 3.29 (similar to a Z-score of 3.29, corresponding to p = 0.001). Only clusters that included at least 30 voxels are reported (k > 30, voxel size 2 x 2 x2 $mm^3$).

| Region | x | y | z | Bootstrapped ratio | k (cluster size) |
|---|---|---|---|---|---|
| **Increased activity** | | | | | |
| Right precuneus | 46 | 26 | 64 | 6.7696 | 4376 |
| Right middle frontal | 55 | 69 | 66 | 8.1482 | 3247 |
| Left middle frontal | 22 | 66 | 66 | 7.4470 | 2348 |
| Left angular | 20 | 29 | 60 | 6.0004 | 1615 |
| Left supplementary motor | 40 | 67 | 62 | 6.1646 | 934 |
| Left cerebellum crus I | 27 | 29 | 19 | 4.6015 | 470 |
| Right cerebellum crus I | 59 | 26 | 19 | 4.8791 | 379 |
| Left middle temporal | 10 | 42 | 30 | 5.6573 | 185 |
| Left cerebellum crus II | 33 | 19 | 23 | 4.3037 | 107 |
| Left middle orbitofrontal | 20 | 90 | 35 | 3.9661 | 71 |
| Right cerebellum crus II | 44 | 22 | 22 | 4.6434 | 58 |
| Right inferior temporal | 71 | 36 | 30 | 4.2383 | 58 |
| **Decreased activity** | | | | | |
| Left superior temporal | 12 | 47 | 42 | -6.0466 | 928 |
| Right superior temporal | 73 | 49 | 41 | -4.5926 | 373 |
| Right superior temporal | 68 | 60 | 38 | -5.0179 | 332 |
| Left precuneus | 35 | 33 | 43 | -4.3832 | 296 |
| Right superior occipital | 53 | 13 | 43 | -4.8425 | 286 |
| Left postcentral | 20 | 47 | 69 | -4.5957 | 224 |
| Right postcentral | 69 | 55 | 61 | -4.6700 | 146 |
| Left superior occipital | 33 | 13 | 46 | -4.8137 | 141 |
| Right superior occipital | 49 | 17 | 53 | -4.1622 | 124 |
| Right superior medial frontal | 41 | 90 | 42 | -3.9361 | 94 |
| Right insula | 59 | 63 | 42 | -4.3637 | 67 |
| Left medial orbitofrontal | 40 | 92 | 32 | -4.4728 | 58 |
| Left supramarginal | 10 | 50 | 58 | -4.3661 | 50 |
| Left superior temporal | 9 | 55 | 33 | -4.2464 | 30 |
